# Comparative proteomic analysis of *Fusarium oxysporum* f. sp. *cubense* strains (*Foc* R1 and *Foc* TR4) provides better insights into mechanisms of their virulence, habitat adaptation and pathogenesis

**DOI:** 10.1101/2021.12.29.474485

**Authors:** Thangavelu Raman, Kalaiponmani Kalaimughilan, Edwin Raj Esack

**Author notes:** Corresponding author:* R. Thangavelu, Plant Pathology Division, ICAR-National Research Centre for Banana, Tiruchirappalli-620 102, Tamil Nadu, India.

## Abstract

*Fusarium oxysporum* f. sp. *cubense* (*Foc*), a devastative soil-borne fungal pathogen causing vascular wilt (i.e. Panama disease) which leads to severe crop losses in most of the banana-growing regions of the world. As there is no single source of effective management practices available so far, understand the pathogenicity of the organism may help in designing effective control measures through molecular approaches. The study aims to compare the proteome of the two pathogenic *Foc* virulent strains, Race 1 (*Foc* R1) and tropical race 4 (*Foc* TR4) that are capable of infecting the Cavendish group of bananas using 2-dimensional (2-D) gel electrophoresis, MALDI-TOF/MS and MS/MS analysis. The results of the study revealed that the proteins, peroxiredoxins, NAD-aldehyde dehydrogenase (NAD-ALDH), MAPK protein, pH-response regulator protein palA/rim-20 and isotrichodermin C15 hydroxylase have shared homology with the fungal proteins, which regulate the osmotic stress response, signal transduction, root colonization and toxin biosynthesis. These are the important functions for the pathogen survival in an unfavourable environment, and successful establishment and infection of the banana host. The present study also identified several putative pathogenicity related proteins in both *Foc* R1 and *Foc* TR4. Specifically, certain *Foc* TR4 specific putative pathogenicity related proteins, phytotoxins biosynthesis gene, fructose 1,6-bisphosphate aldolase class II, Synembryn-like proteins found to contribute strong virulence. Overexpression or knockout of the elective genes could help in devising better control measures for the devastative pathogens in the future. To the best of our knowledge, this is the first report on the proteomics of *Foc* R1 and *Foc* TR4 strains of Indian origin that infect Cavendish bananas.

## Introduction

Banana (*Musa* spp.) is the most important fruit crop in the world where India is the frontrunner with the highest production (FAOSTAT 2018). Though hundreds of banana cultivars have existed globally and in India, Cavendish (AAA) and Plantain (AAB) subgroups registered ∼64% of total banana production (Ploetz 2015). As like any other living organism, the banana plants encounter numerous biotic as well as abiotic challenges in their life cycle. *Fusarium oxysporum* f. sp. *cubense* (*Foc*) is the chief biotic factors that seriously causes wilt disease in banana (i.e. Fusarium wilt) in terms of growth and production. In the severe conditions the disease leading to the death of the infected plant and thus complete yield loss.

Fusarium wilt in banana was first reported during the 18^th^ century in a cv. Gros Michel (AAA) and was identified to be infested by *Foc* Race-1 (*Foc* R1) (2015). In the early 19^th^ century, the devastation was increased and scarcity of *Foc* R1 free soil pushed the Caribbean commercial banana planters to take initiative in search of alternative resistant cultivars. This exploration resulted in the introduction of the Grand Nain bananas (AAA) (Stover 1962), which is resistant against *Foc* R1. As a result, the wilt incidence in the cv. Gros Michel becomes disappeared as a threatening problem of the banana plantations (Buddenhagen 1990). Black leaf streak caused by *Pseudocercospora fijiensis* (anamorph *Mycosphaerella fijiensis*) became the new primary disease of banana and Fusarium wilt was forgotten as a threat until the reemergence of a micro evolved variant of *Foc* R1 i.e. *Foc* tropical race 4 (*Foc* TR4) in the late 19^th^ century (Buddenhagen 1990). The wilt disease by the *Foc* TR4 strain has become even worse during the 21^st^ century in Southeast Asia. However, the emergence, spread, and outbreaks of *Foc* TR4 have been certainly reported from various parts of Africa, India, Pakistan, Thailand, Cambodia, Australia, China, Indonesia, Jordan, Lebanon, Malaysia, Mozambique, Oman, the Philippines, Taiwan, Vietnam and South America (Ploetz 2006; García-Bastidas et al. 2014; Ordoñez et al. 2016). The disease pandemic is due to limited bananas genetic base and speedy spread and/or adaptation of *Foc* TR4 to its given environment, which presents a dangerous situation for global banana production. Thus, the need of the hour is to understand the race physiology of the pathogen and its virulence factors besides the diversification and expansion of the banana genome are the strategies to overcome the disastrous disease. The best efforts and major investments have recently been made to quarantine the epidemics of *Foc* by the researchers across the world through investigating the biology, genetic diversity, pathogenicity and virulence of different *Foc* races using the advanced ‘*omics’* techniques and tools.

Among the ‘*omics’* techniques, proteomics is of particular interest because it is a powerful technique capable of resolving thousands of proteins simultaneously and thereby gives a system-wide view on the molecular response of an organism (Giaever et al. 2002; Grinyer et al. 2004; Marra et al. 2006). The integrated technique involves 2-D gel electrophoresis, mass spectrometry and bioinformatics analyses for the separation, identification and annotation of proteins. Improvements in proteome technologies are helping to understand a large number of proteins in several pathogenic filamentous fungi (Kim et al. 2007). In addition to analyze the gene expression (*i*.*e*. mRNA), it is desirable to study the proteome because highly expressed genes do not necessarily correlate with the protein expression and their turnover in the given time (Tsai-Morris et al. 2010). Moreover, the genes required for a response are not necessarily the same genes that are differentially regulated as a result of the response (Giaever et al. 2002). Spliced variants of a gene are also known to code altogether difference in protein structure (Cheah et al. 2007) as a post-translational modification which may lead to multiple functions of a protein (de la Cadena et al. 2007). Besides, the proteomics study can be used to explore the host-pathogen interactions, which is specific to a location or tissue as well (Rampitsch et al. 2006; Marra et al. 2006). In a wider context, separation and identification of protein related to pathogenicity can provide researchers to knock down the elective candidate/target genes in the fungal genome as a measure of control or to elucidate systems biology (Giaever et al. 2002).

As proteins are directly related to cellular functions and molecular response in the biological system, the differences observed in the protein(ome) identified from 2-D gels of the two contrasting important pathogenic strains will assist in distinguishing unique and shared pathogenicity or virulence factors. Therefore, the main objective of this study is to compare the proteome of *Foc* R1 and *Foc* TR4 that affects the Cavendish group of bananas by 2-D electrophoresis technique, MS/MS and MASCOT interface. Moreover, the proteomics study also identifying the putative virulence factors, which may assist in explaining pathogenicity specific targets that can be used as candidate genes/proteins for the diagnosis of disease and develop management approaches.

## Materials and Methods

### Fungal isolates and culture conditions

Two *Foc* strains, *Foc* R1 (Thangavelu and Mustaffa 2010) and *Foc* TR4 (Thangavelu et al. 2019) used in the present study were availed from the Plant Pathology Division, ICAR-National Research Center for Banana *Foc* culture collection. Seven-days-old mycelium of the single spore culture of *Foc* strains grown on potato dextrose agar (PDA) medium at 25±2 °C was further transferred to 150 mL potato dextrose broth medium and incubated for seven days at 25±2 °C under constant shaking of 120 rpm. The mycelia were filtered through cheesecloth and washed with ice-cold 0.1 mM Phenyl methyl sulfonyl fluoride. Mycelia were then blotted dry with sterile filter paper stacks and used immediately for protein extraction.

### Protein extraction

Protein extraction from air-dried *Foc* mycelium was performed according to Kalaiponmani et al (Kalaiponmani et al. 2017). Briefly, 500 mg of fine mycelial powder was homogenized in ice-cold TCA-Acetone buffer (20% TCA in ice-cold acetone, 0.2% DTT) for 5 min and precipitated overnight at -20 °C. The samples were then centrifuged at 12,000 rpm for 30 min at 4 °C. The collected pellet was washed two times with wash buffer (ice-cold acetone, 0.2% DTT). The pellet was then completely dissolved in 1.2 mL of extraction buffer (1:1 phenol pH 8, 30% (w/v) sucrose, 2% (w/v) SDS, 5% *β*-mercaptoethanol and 0.1 M Tris-HCl) and incubated at 4 °C for 5 min. Following centrifugation at 12,000 rpm for 5 min, the upper phenol phase was transferred into a new tube and protein was precipitated with 0.1 M ammonium acetate in 100% (v/v) ice-cold methanol for 2 h at -80 °C. The white pellet was obtained by centrifugation at 12,000 rpm for 15 min at 4 °C and washed twice with methanol (100%) and once with acetone (80%). The pellet was then air-dried at room temperature and dissolved in rehydration buffer (7 M urea, 2 M thiourea, 4% CHAPS, 0.8% IPG-buffer, 20 mM DTT). Protein concentration was estimated according to the Bradford method (Bradford 1976) against Bovine Serum Albumin as standard and extracted protein was stored at -20 °C until further use.

### 2-D gel electrophoresis

For the first dimensional isoelectric point-based separation, 250 μg of total protein was loaded on a 13 cm Immobiline dry-strip (pH 4-7, M/s. GE Healthcare Life Sciences, Amersham, UK), on a rehydration tray. The strip was covered with white mineral oil (DryStrip Fluid, M/s. GE Healthcare Life Sciences, Amersham, UK), and incubated horizontally at 22 °C for 18 h. Protein focusing to its corresponding isoelectric point within the strip was achieved by following three-step protocol: 500 V for 1 h (linear), 1000 V for 1 h (linear), and 3500 V for 28 kVh (linear) with a Hoefer^™^ IEF 100 (M/s. GE Healthcare Life Sciences, Amersham, UK) at 20 °C. For the second dimensional molecular weight-based separation, the strips were equilibrated with 10 mL of equilibration buffer I (50 mM Tris–HCl, 6 M urea, 30% (v/v) glycerol, 2% (w/v) SDS, 0.002% (v/v) bromophenol blue) for 12 min containing 1% (w/v) dithiothreitol. A second equilibration was then performed with 2.5% (w/v) iodoacetamide added to the SDS equilibration buffer. The equilibrated strips were placed on lab cast 1.5 mm SDS-PAGE gels (12.5%) and the strips were sealed with 1% (w/v) agarose solution containing tracking dye and run on a vertical electrophoresis unit (Hoefer^™^ SE600 Ruby) at 10 mA per gel. For each sample, duplicate gels were electrophoresed on two separate occasions.

### Visualization and image analysis

Electrophoresed gels were visualized by colloidal Coomassie Brilliant Blue (CBB) staining (Neuhoff et al. 1988). Briefly, the gels were fixed for 2 h in a fixing solution (40% (v/v) methanol and 7% (v/v) glacial acetic acid), washed twice with double distilled water for 20 min and stained overnight in a staining solution (10% (v/v) methanol, 0.1% (w/v) CBB G-250, 1.6% (v/v) O-phosphoric acid, and 10% (w/v) ammonium sulphate). Stained and washed gels were scanned using EPSON^®^ Perfection 750 Pro Scanner (EPSON^®^ Inc. USA). Careful inspection and comparison of the normalized proteome profiles from the two isolates shown qualitative and quantitative variations. Spot detection, measurement and matching were done using Hoefer^™^ 2-D view software (Non-linear, USA, Inc.) according to the standard protocol. Three biological replicates were analyzed for each sample. Overall, protein spots were considered to be differentially expressed only if at least of twofold intensity variations were detected and only those spots were subjected to further analysis.

### MS/MS (tandem mass spectrometry) analysis and database search

Selectively excised spots from the preparative gels (stained with CBB) were sent to the proteomics facility, Molecular Biophysics Unit, Indian Institute of Science, Bengaluru, India, for MALDI-TOF/MS and MS/MS analysis. The resulted mass fingerprint and fragmentation spectra data of peptides were searched against taxonomy fungi in NCBI entries using MASCOT server (www.matrixscience.com) to classify the functions of the peptide fingerprints with the following parameters: one missed cleavage site, 100 ppm mass tolerance, carbamidomethyl modifications of the cysteine’s as a fixed modification and methionine oxidation as a variable modification. The four criteria *viz*., significant score, percentage of sequence coverage, E-value of the homologous protein and minimum of five-matched peptides were followed to increase the confidence in the protein identification. Based on the function (Blast2Go) and putative function of the protein spot assigned, the relevance of the pathogenicity of the respective strains is discussed.

## Results and Discussion

### 2-D protein profile of Foc global proteome

The global (total) mycelial proteome of two contrasting Indian *Foc* races that are capable of infecting Cavendish bananas was established using IPG strip covering the pH range of 4-7. In our experimental conditions, ∼600 spots were recognized in both the strain by 2-D image analysis (see Fig. 1 and 2). The combined image analysis revealed that there were 82 protein spots that significantly (p<0.05) exhibited more than a twofold spot intensity in both of the strains. Among the elective protein spots eluted from the gel, MS/MS and MASCOT interface based database search has identified 46 protein spots from both the isolates, which are indicated by Arabic numerals on the gels (Fig. 1 and 2). Out of 46 spots, 25 are from *Foc* R1, and 21 are from *Foc* TR4 (see Supplementary Figure-1).

**Figure 1.**
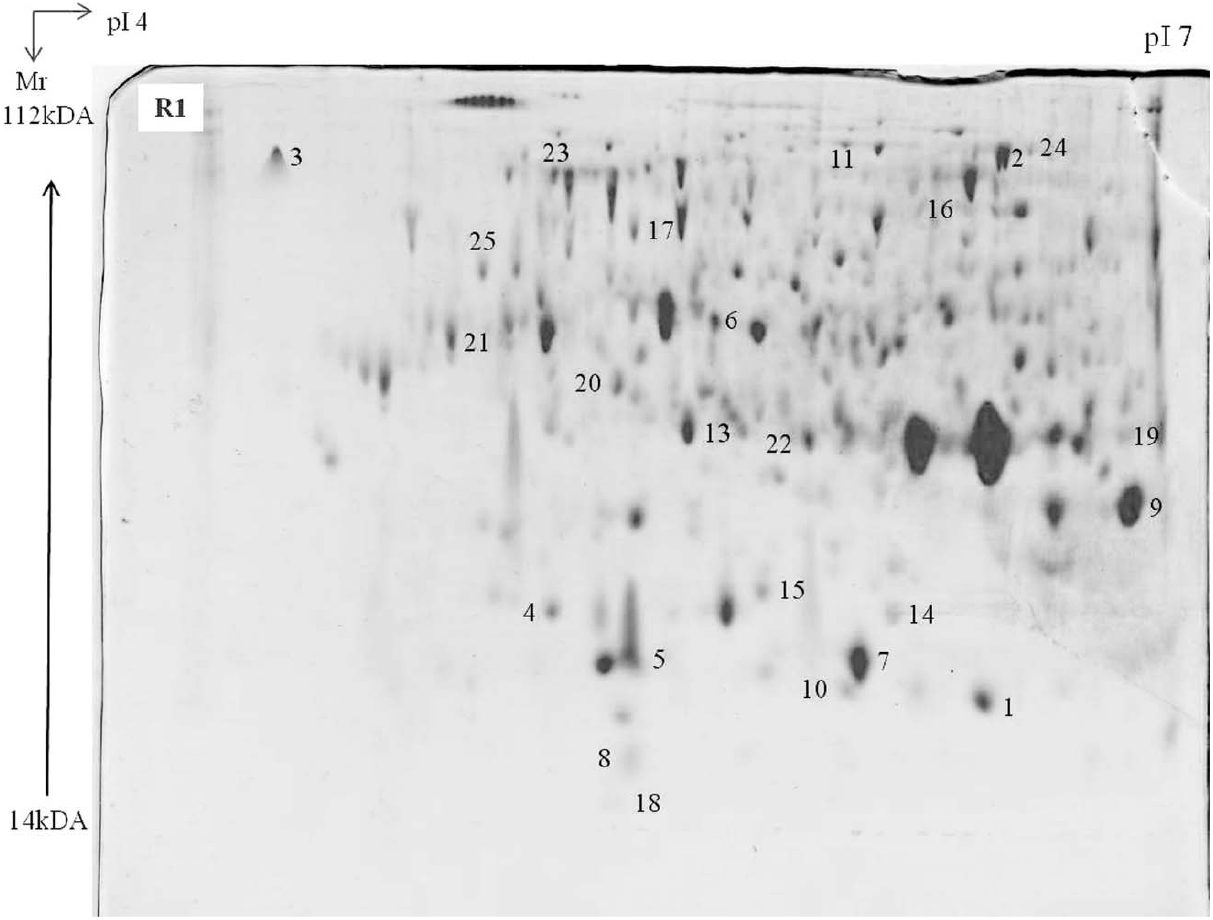
Proteome map of *Fusarium oxysporum* f. sp. *cubense* race 1 (R1).

**Figure 2.**
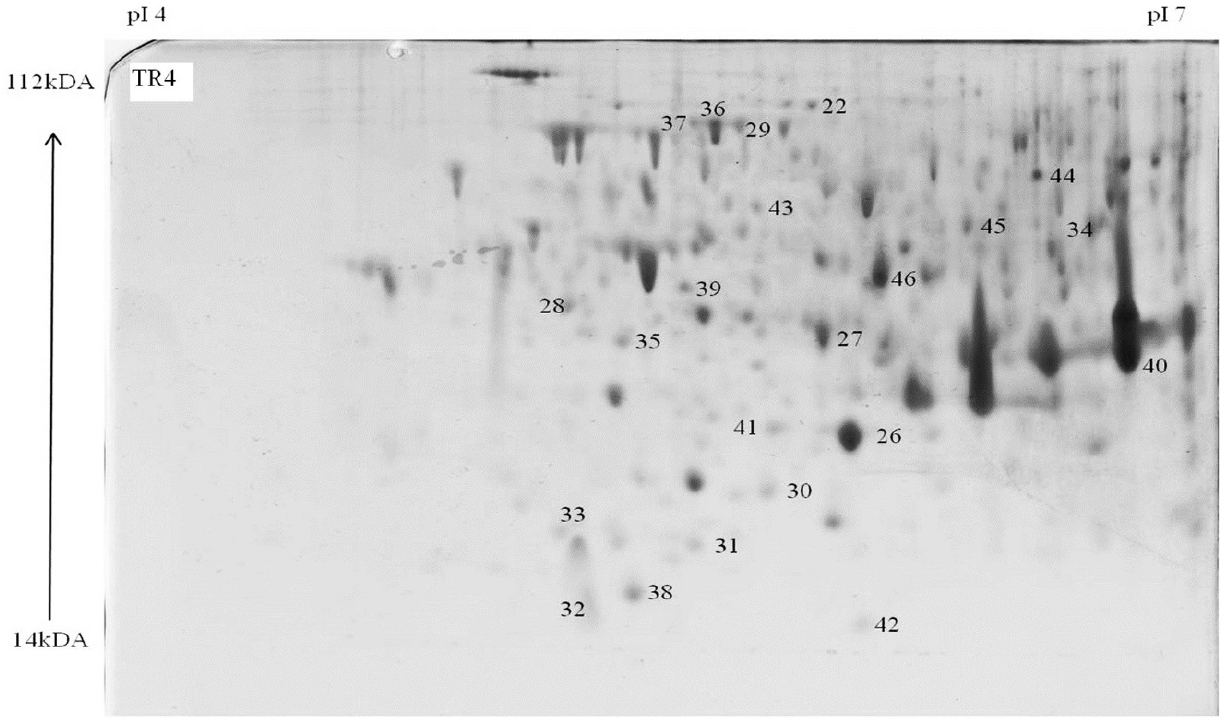
Proteome map of *Fusarium oxysporum* f. sp. *cubense* tropical race 4 (TR4).

Based on the global MASCOT and BLASTp annotations, most of the proteins were homologous to fungal proteins, which are involved in cell wall degradation, osmotic stress response, signal transduction and homeostasis, ATP synthesis and conservation, amino acid transport and metabolism, biosynthesis of secondary metabolites, and proteins & lipids transport and metabolism. The percent coverage of amino acid sequence, number of matched peptides, the score of the identified proteins, accession number and spot annotation description are presented in table-1.

**Table 1:**
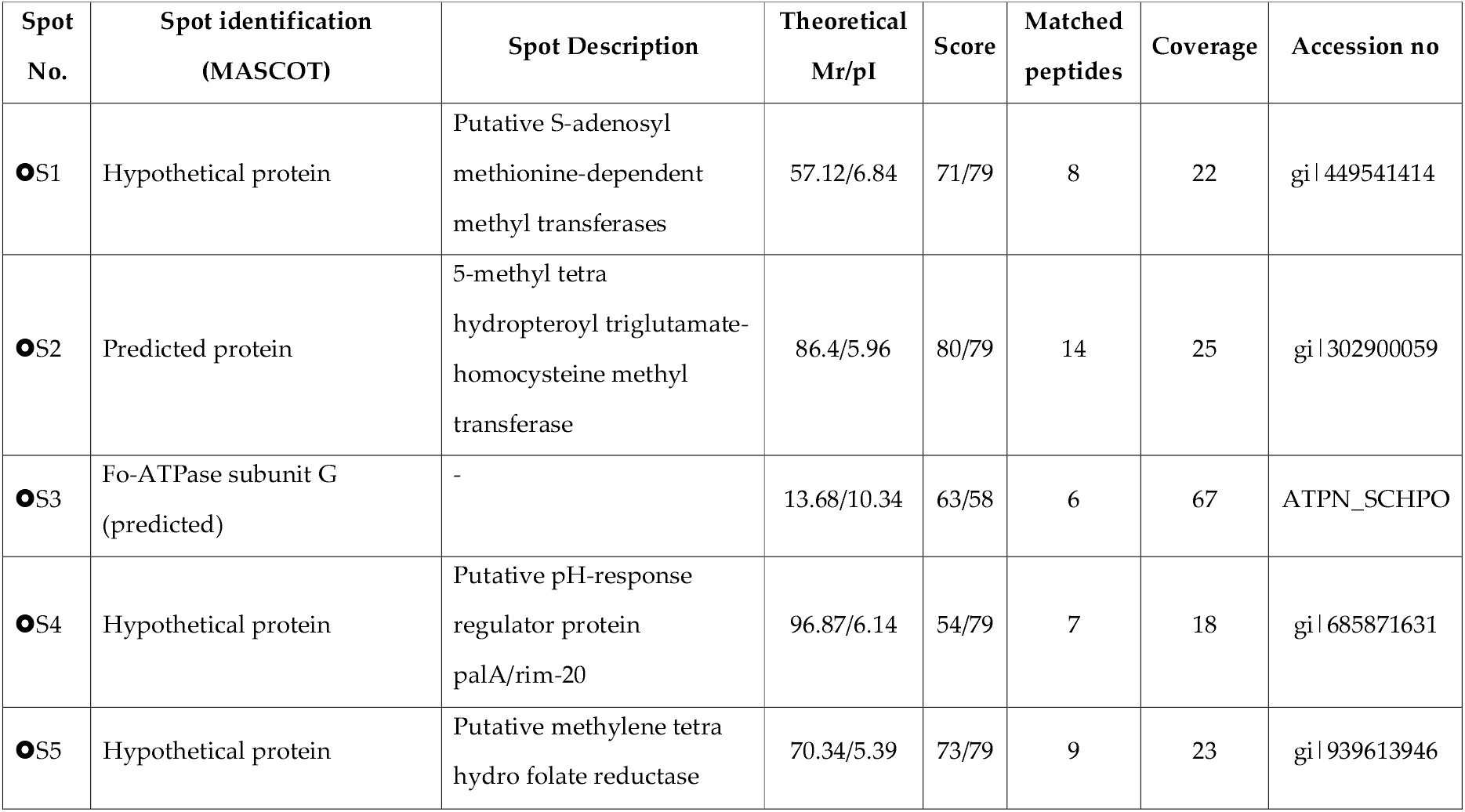
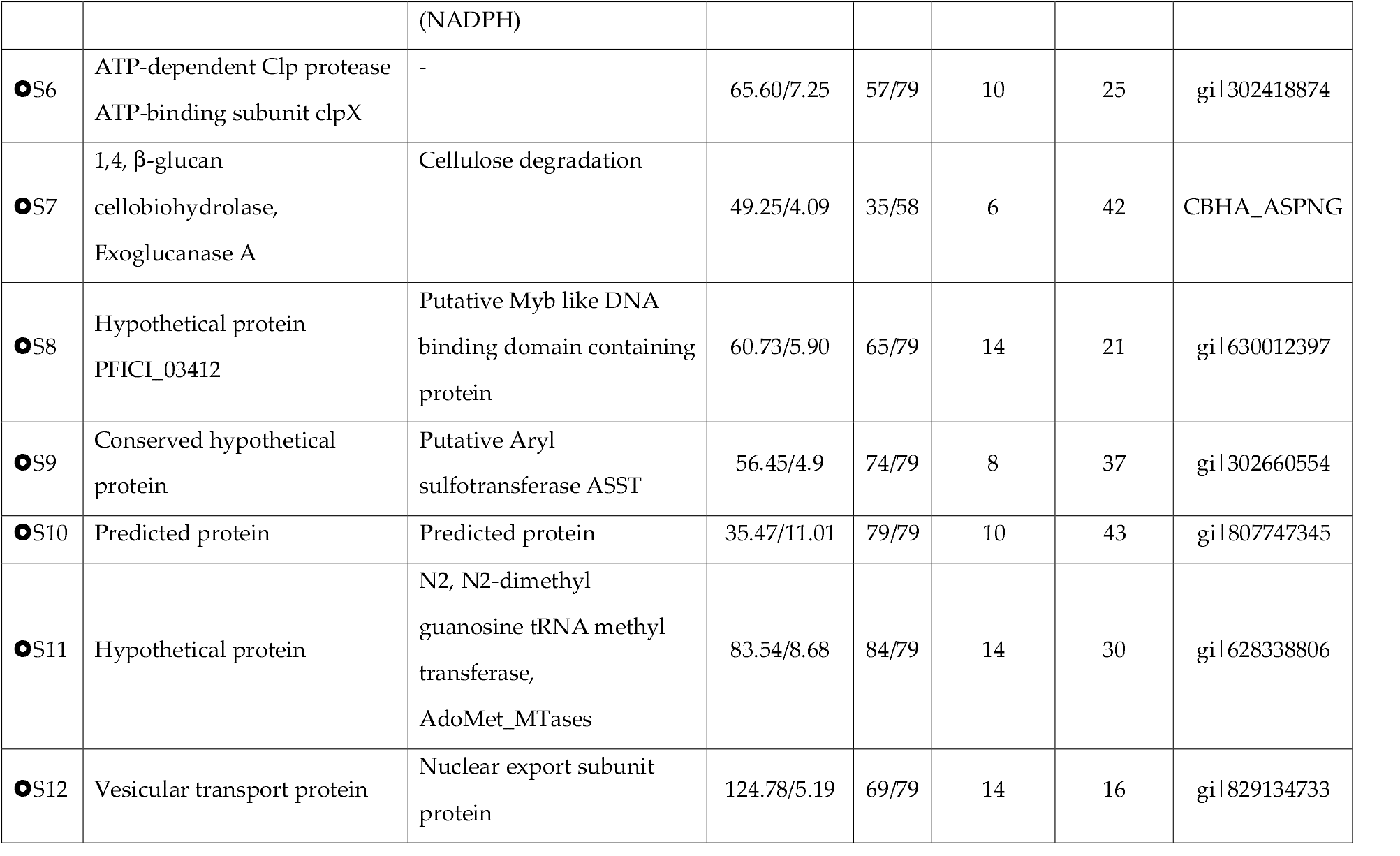
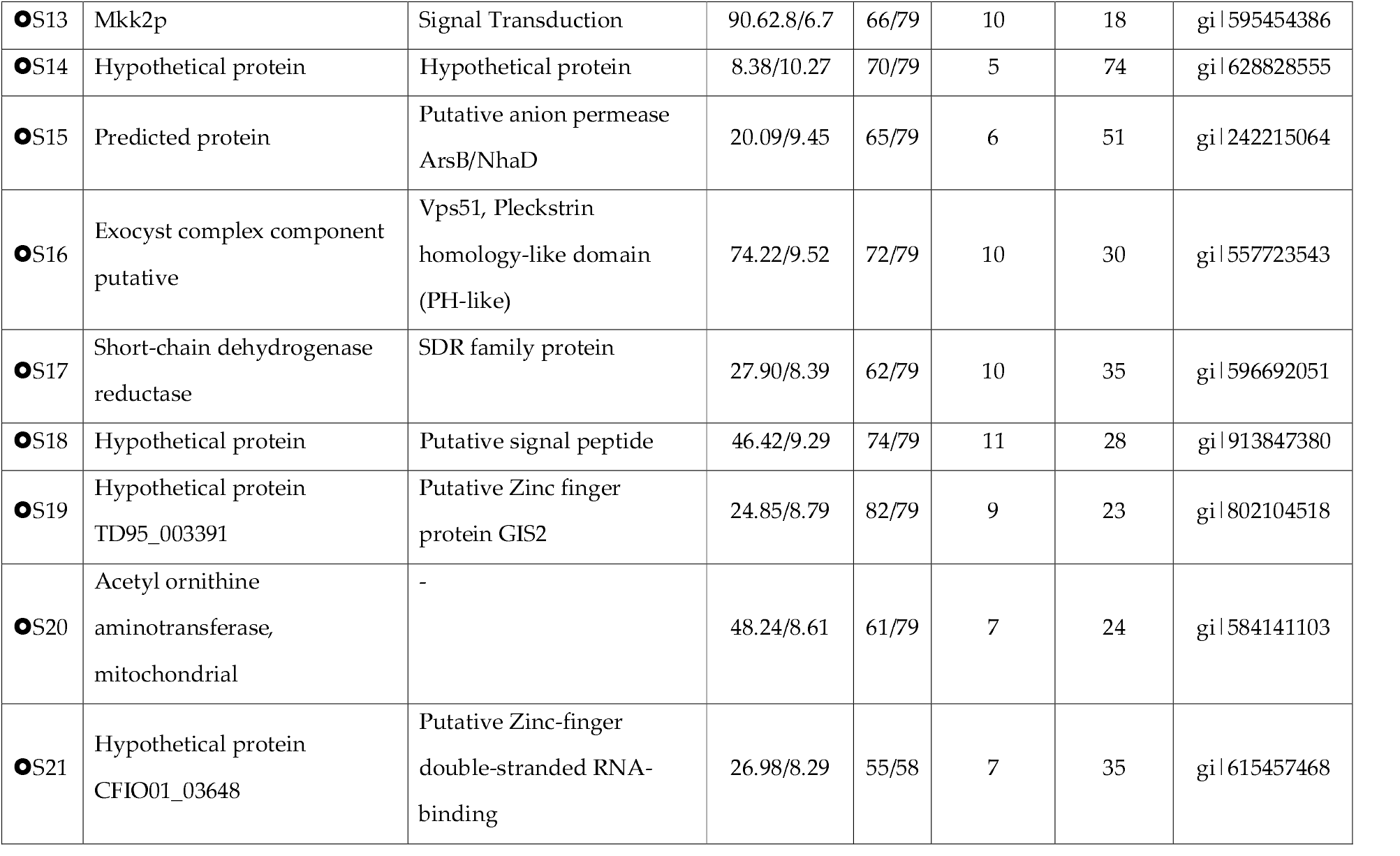
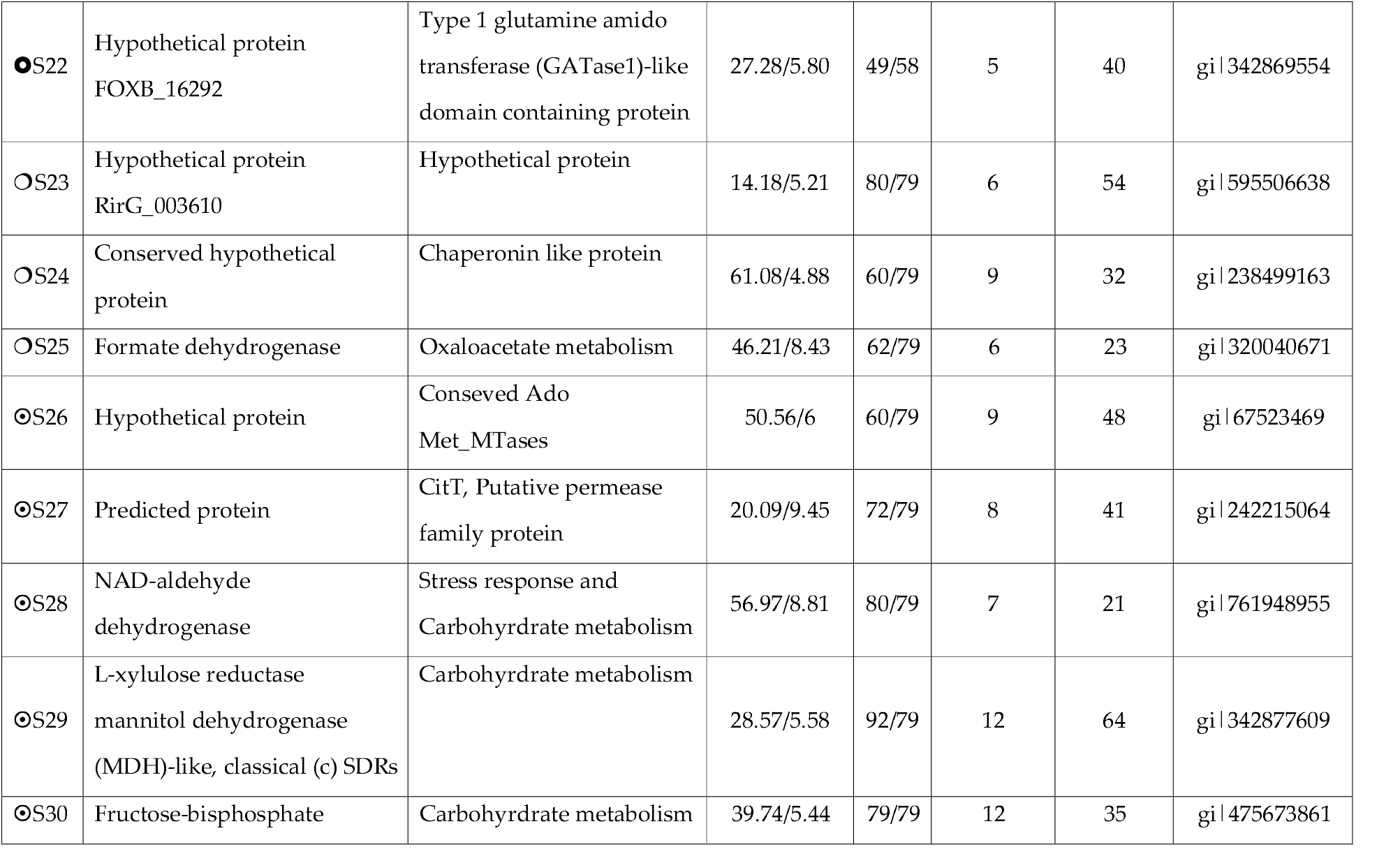
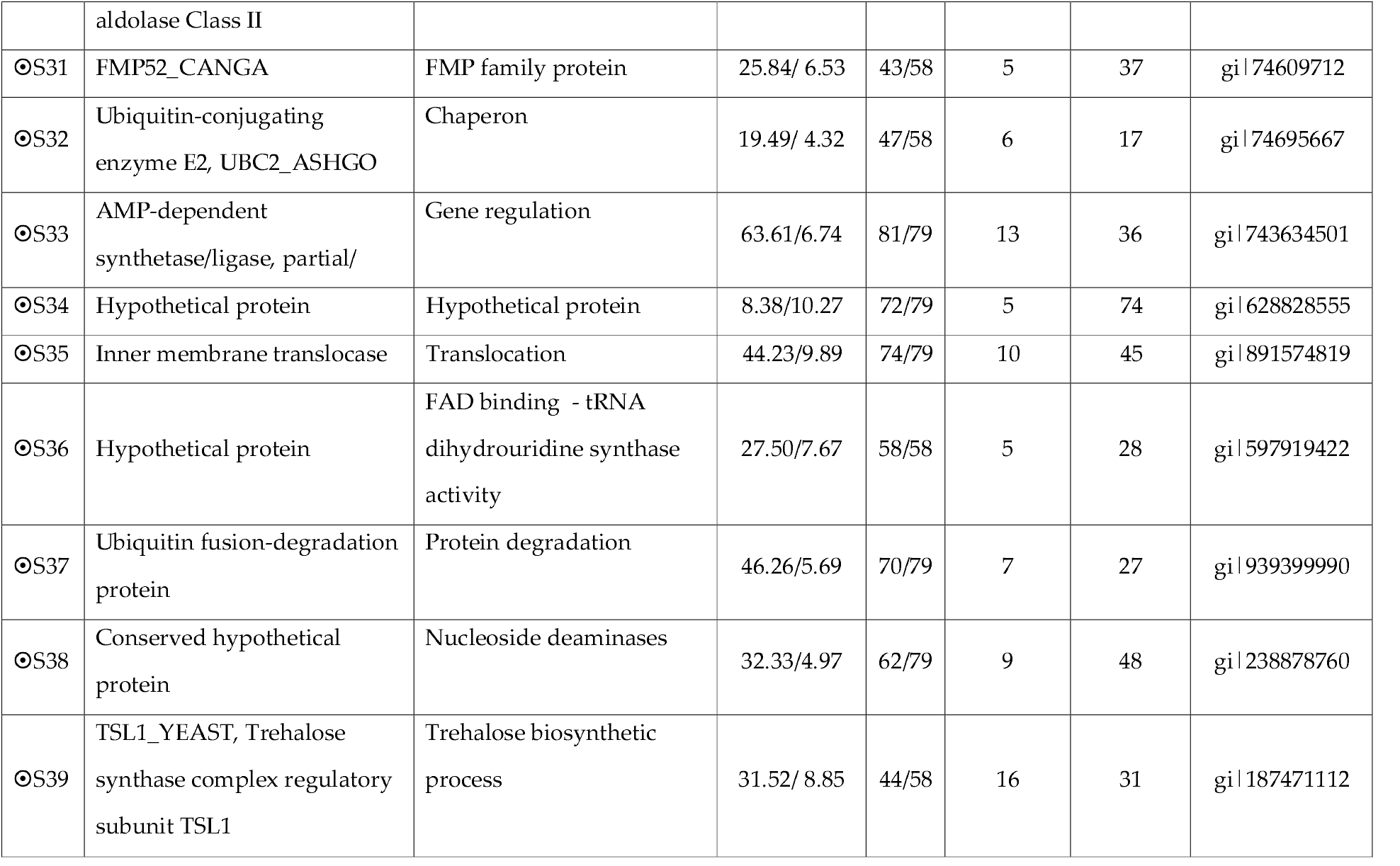
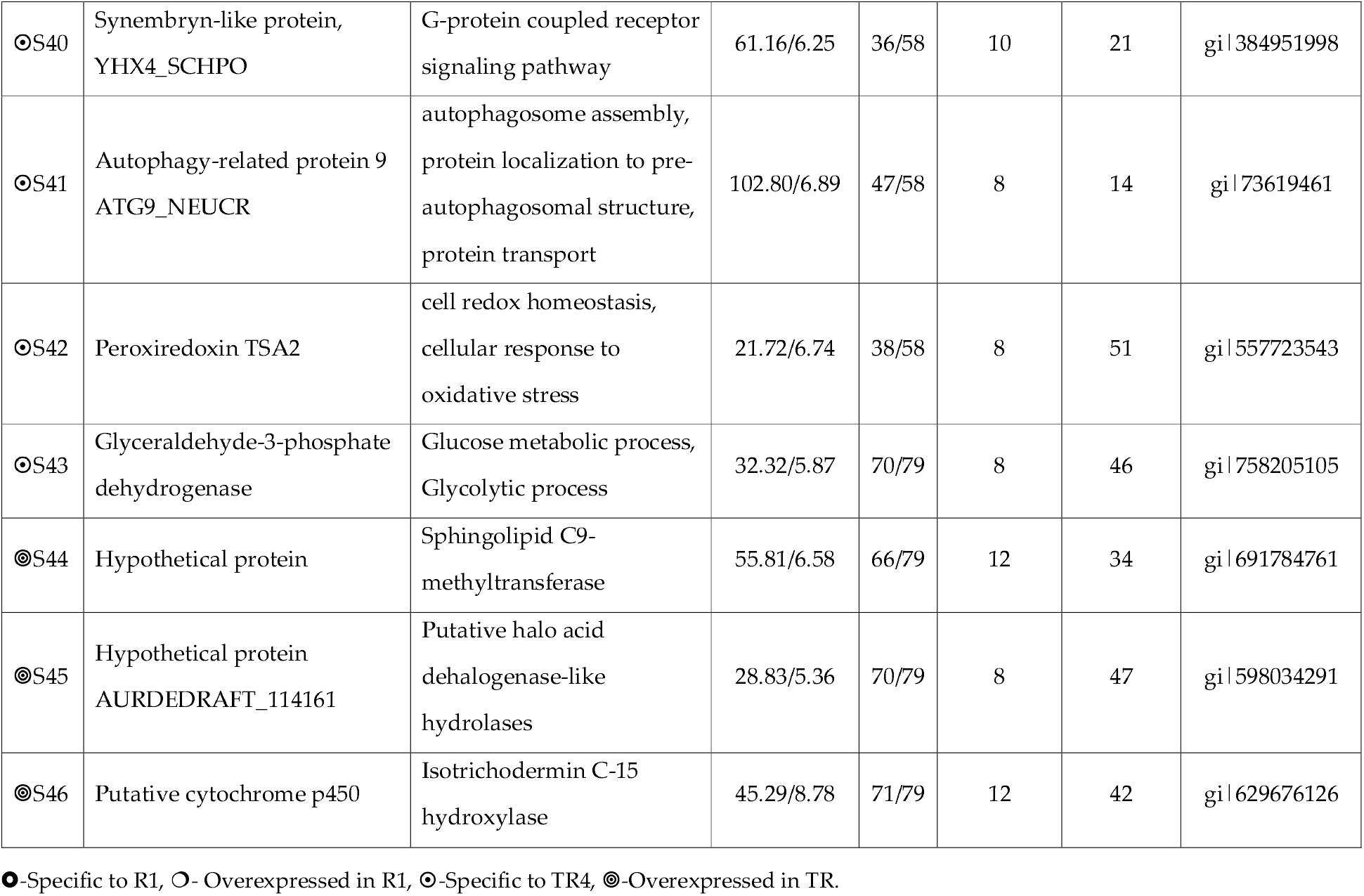
Identification of proteins from *Foc* R1 and *Foc* TR4 isolates of *Fusarium oxysporum* f. sp. *cubense*

### Functional annotation and grouping of proteins

Firstly, a BLASTp search was done to annotate the proteins using the Universal Protein KnowledgeBase (UniProtKB). Secondly, a local search against EuKaryotic Orthologous Groups (KOG) database was carried out. Combining the results of both analyses, the annotated proteins are then categorized into members of the known pathways. KOG based categorization revealed that carbohydrate transport and metabolism-related proteins (4) were most abundant, followed by post-translational modification chaperone related proteins (PTM, 3), lipid transport and metabolism (3), amino acid transport and metabolism (ATM, 3) and translation, ribosomal structure and biogenesis (3) (Fig 3a). The remaining proteins (30) featured various other functional annotations and could not be accommodated in any KOG group. Further, gene ontology (GO) annotation at three different levels namely *viz*., biological processes, cellular components and molecular functions were performed. The GO annotation for biological processes revealed that the most proteins are multifunctional and mainly involved in the metabolic process (26 proteins; GO:0008152) followed by a single organism process (20 proteins; GO:0008150) and cellular process (14 proteins; GO:0009987) (Fig. 3b). The GO annotation for molecular functions showed that the largest portion of proteins has catalytic activity (23 proteins; GO:0003824), followed by metal ion binding (17 proteins; GO:0046872) (Fig. 3c). Finally, the cellular component GO annotation revealed that the largest portion of proteins (14 proteins; GO:0005575) belonged to the cell followed by organelles (11 proteins; GO:0043226) (Fig. 3d). Based on this GO annotation analysis, it can be assumed that *Foc* has a repertoire of protein that belongs to a cell wall with catalytic and ion binding activity. This is an important feature for the quick adaptation of *Foc* to its microclimate and also could reprogramme its whole system according to the external stimuli and strengthen its defense against host response and effective infection (Sun et al. 2014).

**Figure 3.**
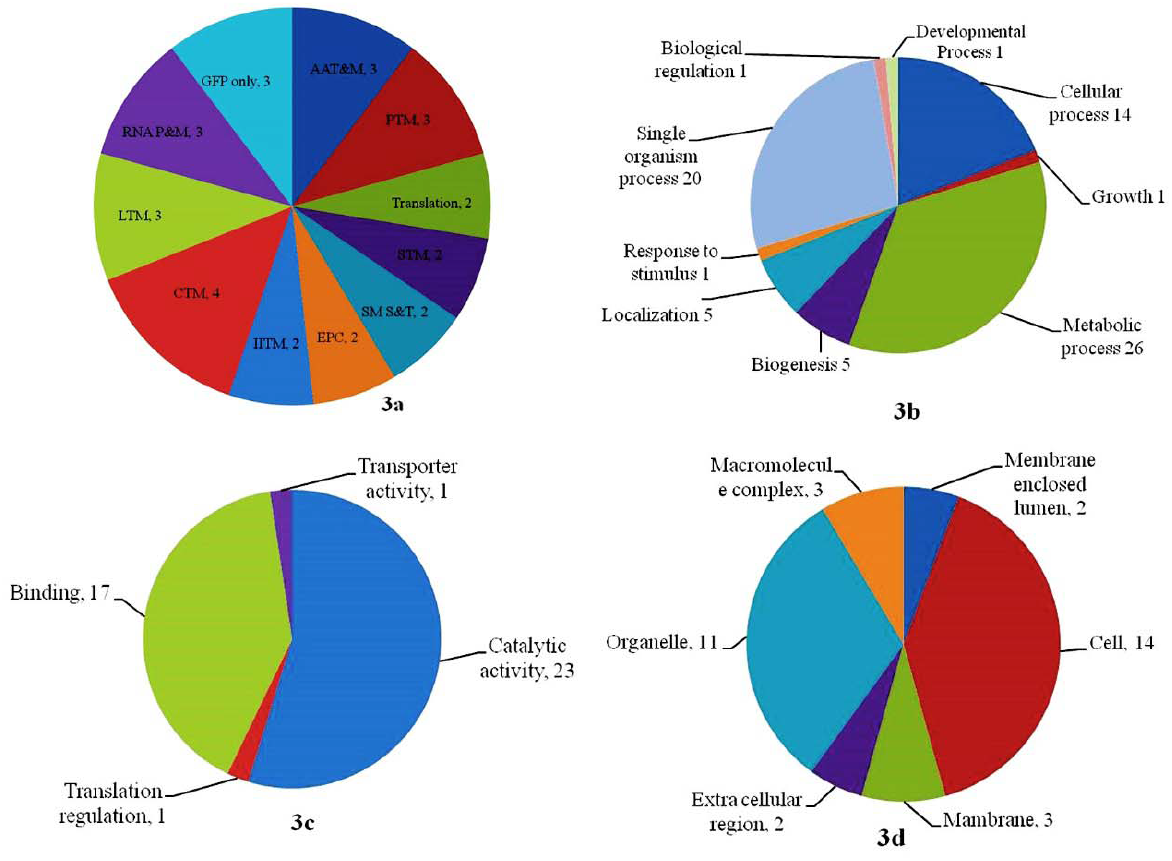
Functional classification analysis of the identified proteins: (3a) the proportion of the identified proteins in each category of the COG classification system, Gene Ontology (GO) based protein classification for biological process (3b), molecular function (3c) and cellular component (3d).

### Differentially expressed proteins in Foc R1

This study recognized 25 protein spots that significantly exhibited more than a twofold spot intensity in *Foc* R1. Comparison of *Foc* R1 with *Foc* TR4 proteome reveals that 22 out of 25 proteins are unique to *Foc* R1 and the remaining three are abundant with *Foc* TR4.

When a pathogen encounters a potential host, the cell wall is the first barrier used to mess with it. However, pathogen tries to colonize the host and derive the energy supplies from the cell wall matrix that comprising with a varying proportion of pectin, hemicellulose, lignin, and structural proteins. The pathogens producing a mixture of enzymes, such as endoglucanases (EGs), exo-glucanases including cellobiohydrolases and β-glucosidases which act on cellulose, xylan, and pectin, and deconstruct them to make an entry into the host cell and derive carbon/food from the disintegrated molecules (Lynd et al. 2008). The 1,4,*β*-glucan cellobiohydrolase of exoglucanase A identified (Spot 7) in the *Foc* R1 may help in the deconstruction of cellulose of the host cell wall during the colonization, infection and disease development or late stages of invasion as like *Botrytis cinerea* (Brito et al. 2006).

Mitogen-activated protein kinase (MAPK) cascades involving in phosphorylating various cellular proteins, which includes transcription factors and other regulatory proteins. The experimental results suggested that silencing of MAPK in *Fusarium oxysporum* led to a reduction in putative pathogenicity functions, such as hyphal growth at the liquid-air interface, colonization of the host and cell wall degrading enzyme, pectate lyase (Di Pietro et al. 2004). Similarly, a null mutant of *F. oxysporum* has led to a substantial reduction in fungal virulence against banana plants (Ding et al. 2015). Therefore, it is assumed that the presence of MAPK protein (Spot 13) in *Foc* R1 is contributing to the virulence and pathogenicity. As like many other soil fungi, *F. oxysporum* also performing anaerobic metabolism of nitrate to form ammonium (Zhou et al. 2002) through (N_2_O) dissimilatory means which support anoxic growth of fungi and modulation of ambient pH to its support (Uchimura et al. 2002). Therefore, formate dehydrogenase (Spot 25) identified in the *Foc* R1 found to enhance the denitrification in a dissimilatory manner and alter the ambient pH to complement the initial infection aspects.

A hypothetical protein of Spot 4 in *Foc* R1 was annotated as a putative pH-response regulator protein, palA/rim-20, which is one of the most specialized pH response pathways directly or indirectly involved in infection, colonization and establishment through proteolytic activation of a zinc finger transcription factor, *pacC* (Penalva and Arst 2002). As *Foc* is soil-born, the presence of a palA/rim-20 in *Foc* R1 revealed that the fungi have strong sensory processes in responding to its microenvironment changes by regulating gene expression accordingly to secrete arsenal of pH manipulating molecules for achieving ambient pH to complete the infection process effectively.

Protein spot 15 was identified as putative anion permease (ArsB/NhaD). These permeases have been shown to translocate sodium, arsenate, antimonite, sulphate and organic anions across biological membranes help in nutrient absorption.

### Differentially expressed proteins in Foc TR4

Among the total mycelial proteome of *Foc* TR4, 21 protein spots exhibited more than a twofold spot intensity, which is subjected to MS/MS analysis and annotated further. Proteome comparison of *Foc* races reveals that 18 out of 21 proteins are unique to *Foc* TR4 and the remaining three are abundant with *Foc* R1.

Several factors ranging from various macro and micro molecules to physical structures that determine the virulence of a fungal pathogen. *Foc* TR4, a proven aggressive strain among the other *Focs* presumed to be dependent upon several pathogenic factors within the plant host (for penetration, survival, growth) as well as environmental factors (i.e. temperature, moisture, light, aeration, nutrient availability and pH) (Alkan et al. 2013). All these factors independently or in combination perturb the fungal cell homeostasis and cause cell death due to excess production of reactive oxygen species (ROS) during lipid peroxidation (Heller and Tudzynski 2011). In this study, three protein spots, namely NAD-aldehyde dehydrogenase, (NAD-ALDH, Spot 28), synembryn-like protein (Spot 40) and peroxiredoxin (Prxs; Spot 42) which are identified to be involved in a cell redox homeostasis and G-protein coupled receptor-signalling pathway besides toxin biosynthesis and cell wall degradation. The presence of *Foc* TR4 specific proteins helps the organism to overcome the host immune response during the invasion, which in turn leads to the completion of the infection process (Cabiscol et al. 2000).

ALDH, an enzyme of ROS detoxification (Jakoby and Ziegler 1990) which provides tolerance to Yeast against to the variety of abiotic and biotic stresses, specifically, osmotic and oxidative stresses and during depletion of glucose reserve (Singh et al. 2013). Hence, the unique presence of NAD-ALDH in *Foc* TR4 implicates that the enzyme protects the organism from osmotic and oxidative stress and even from the depletion of energy source (glucose) during the infection process by activating detoxification of intermediate or exogenous aldehydes. For the successful colonization and infection of the host, pathogenic fungi produce a ROS scavenging ubiquitous enzyme, Prxs (Kim et al. 1988) to counteract the plant defense mechanisms in coordination with the catalase and glutathione peroxidase (Heller and Tudzynski 2011). The presence of Spot 42 in the hypervirulence *Foc* TR4 implies that the pathogenicity determinant, Prxs acts as a ROS scavenger and protectant against the host induced oxidative damage.

The fungal cell highly depends on carbohydrate as a primary source of energy. However, the study identified certain proteins that are involved in alternate carbohydrate metabolic pathways. The enzymes identified in the present study such as L-xylulose reductase (LXR; Spot 29), fructose 1,6-bisphosphate aldolase Class II (FBA-II; Spot 30), trehalose biosynthetic process (TPS3; Spot 39) and glyceraldehyde-3-phosphate dehydrogenase (GAPDH; Spot 43) found to involve in the energy metabolism other than a common carbon source.

Due to short supply of regular carbon sources during the invasion, pathogens acquired energy from alternative carbon sources for the survival, such as amino acids, carboxylic acids (gluconeogenesis and the glyoxylate cycle) (Ramírez and Lorenz 2007). LXR is a short-chain dehydrogenase and reductases superfamily (Kallberg et al. 2002) involved in the assimilation of L-arabinose (Verho et al. 2004), derived from the degradation of cell walls of the banana root (Mojzita et al. 2010), into the fungal pentose phosphate pathway (Klaubauf et al. 2013). FBA-II is a metal-based enzyme present mainly in bacteria and fungi, reversibly catalyzes the cleavage of Fructose 1,6-bisphosphate into triose phosphates (Rodaki et al. 2006). The activation of trehalose pathway is due to the presence TPS3 (Spot 39), which is important for successful infection and survival of the pathogen in the host because the disaccharide trehalose is an energy source as well as osmotic stress protectant during the infection process (Petzold et al. 2006). GAPDH (Spot 43) is localized at the cell surface and play major roles in host-pathogen interactions specifically binding the conidia to a host (Broetto et al. 2010), besides, biosynthesis of volatile sesquiterpenes and catalyze the sixth step of glycolysis (Pachauri et al. 2019).

Considering from the above facts, the presence of GAPDH, FBA-II, LXR, and TPS3 in the *Foc* TR4 which enabling them to consume sugars both carbon acquired from the cytosol of the host cell and carbon derived from the cell wall by degradation. Based on the results of (Yin et al. 2015), (Broetto et al. 2010) and (Petzold et al. 2006), we perceive the following mechanisms by which the hypervirulent strain of Indian *Foc* TR4 has successfully infest and devastate the host are: GAPDH facilitate binding of conidia, enables colonization and establish the interaction with the host; FBA-II is weakening the host cell by exhausting energy through reverse utilization of host cell fructose sugar; LXR enables degradation of the host cell wall and actives L-arabinose assimilation pathway; TPS3 makes trehalose as an alternative energy source and osmotic stress protectant against host ROCs. We also hypothesize that if we can able engineer host-induced silencing of FBA-II gene in the *Foc* TR4 it could be a promising molecular disease control measure against the *Foc* TR4 in the future.

The result of the total mycelial proteome of the Indian *Foc* TR4 revealed the presence of the potent natural mycotoxin biosynthetic pathway of trichothecenes mediated by isotrichodermin C15 hydroxylase (Abbas et al. 1991; Jin 1996). The toxin inhibiting the physiological processes in the host cell and its surrounding points of infection, enabling the spread of the disease (Feys and Parker 2000). Hence, the identified isotrichodermin C15 hydroxylase enzyme in this study (Spot 44) implicated to be involved in C15 hydroxylation of trichothecene biosynthesis, which in turn acts as a potent phytotoxin to banana either in the early phase of infection or in the later stages of wilt disease development (Alexander et al. 1998).

Inter-intra cellular communication and environment sensing are crucial for the survival of the pathogen in the host in which many signaling and messenger molecules and cell surface receptor sensors are involved. G protein-coupled receptors (GPCRs; Spot 40), the largest class of plasma-membrane-localized protein receptors acts as a liaison agent between the host and pathogens, and play an essential role in pathogenicity (Kroeze 2003) through nutrient sensing, pheromone response, fruiting body development and pathogenesis in coordination with intracellular heterotrimeric G proteins (Truesdell et al. 2000; Jain et al. 2002; Grinyer et al. 2004; Yamagishi et al. 2006; Guo et al. 2016). The conserved ado-met methyltransferase (MTases; Spot 26) is a member of the methyltransferases family that act on a wide range of substrates and maintains the cellular homeostasis through sensing external stimuli, adaptation to new conditions, detoxification of toxic molecules (Bedford and Richard 2005) and responding to external perturbation (Springer et al. 1979). From the above facts, it is apparent that the presence of GPCRs and MTases enzymes in *Foc* TR4 may help in nutrient sensing, fruiting body development and pathogenesis besides to adapt to its microenvironment along with other such factors.

Other identified proteins involved in cellular metabolic activities that regulate the protein synthesis and modifications are Ubiquitin-conjugating enzyme (Spot 32), Ubiquitin fusion-degradation protein (Spot 37) and nucleoside deaminases (Spot 38) in the regulation of nucleosides.

## Conclusion

Advancements in “*Omics*”, especially proteomics study opened many doors to understand the pathogenicity, virulence and its associated factors of the plant pathogenic filamentous fungi. In the present investigation, we detected various potential pathogenicity or virulence factors expressed in the *Foc* R1 and Foc TR4 strains, which will help us to identify the factors that are messing with the banana plant and design target-directed tool for Fusarium wilt control. Several proteins identified in this study are involved in various pathogenesis-related functions like osmotic stress response and homeostasis, cell wall degradation, alternate carbon source metabolism, signal transduction, colonization, infection establishment and phytotoxins synthesis and there may be regarded as potential pathogenicity factors. Most of the putative pathogenicity factors were differentially expressed in *Foc* TR4 strain, which supports their role in the aggressive nature of pathogenesis. Few potential putative pathogenicity factors identified, such as the enzymes involved in phytotoxins biosynthesis, alternate carbon source metabolism FBA II and cell signaling components like G protein-coupled receptors shall be utilized as a target for host induced gene silencing and resistance improvement. In the future, in addition to validation of putative pathogenicity factors identified, more comprehensive proteomics of *Foc*-host interactions combined with functional genomics studies using the same model would be useful in elucidating the basis of the molecular interactions in this pathosystem. To the best of our knowledge, this is the first report on the proteomics of *Foc* R1 and *Foc* TR4 strains of Indian origin.

## Acknowledgements

The authors wish to acknowledge the financial assistance of the Indian Council of Agricultural Research (ICAR) to carry out this research work.

## Author Contributions

RT: Outlined the experiment, inspected the entire study, created two-dimensional electrophoresis facilities for the study and written manuscript; KK: Performed the laboratory experiments, data analysis and drafted the manuscript; and ER: reviewed the results and written the manuscript. The manuscript was written through the contributions of all authors. All authors have approved the final version of the manuscript. All the authors have contributed equally.

## Ethical approval

This article does not contain any studies with human participants or animals performed by any of the authors.

## Data Availability Statement

The data sets supporting the conclusions of this article are included. The datasets generated and analysed during the current study are available in the Mendeley repository under the DOI number 10.17632/bkctm5wy8w.1.(Thangavelu et al. 2020)

